# Cognitive Phenotype Shifts in Risk-Taking: Interplay of Non-Suicidal Self-Injury Behaviors and Intensified Depression

**DOI:** 10.1101/2023.12.10.570987

**Authors:** Yi-Long Lu, Yuqi Ge, Mingzhu Li, Shutian Liang, Xiaoxi Zhang, Yupeng Sui, Lei Yang, Xueni Li, Yuyanan Zhang, Weihua Yue, Hang Zhang, Hao Yan

**Author notes:** corresponding authors. (H.Y.), (H.Z.). co-first authors (Yi-Long Lu, Yuqi Ge).

## Abstract

**Background:** Non-suicidal self-injury (NSSI) behavior is significantly prevalent in both adolescents and psychiatric populations, particularly in individuals with major depressive disorder (MDD). NSSI can be considered a result of risky decision-making in response to negative emotions, where individuals choose self-harm over other less harmful alternatives, suggesting a potential decision-making deficit in those engaging in NSSI. This study delves into the complex relationship between NSSI and depression severity in decision-making and its cognitive underpinnings.

**Methods:** We assessed decision behaviors in 57 MDD patients with NSSI, 42 MDD patients without NSSI and 142 healthy controls using the Balloon Analogue Risk Task, which involves risk-taking, learning, and exploration in uncertain scenarios. Using computational modeling, we dissected the nuanced cognitive dimensions influencing decision behaviors. A novel statistical method was developed to elucidate the interaction effects between NSSI and depression severity.

**Results:** Contrary to common perceptions, we found that individuals with NSSI behaviors were typically more risk-averse. Meanwhile, there was a complex interaction between NSSI and depression severity in shaping risk-taking behaviors. As depressive symptoms intensified, these individuals with NSSI began to perceive less risk and behave more randomly.

**Conclusions:** This research provides new insights into the cognitive aspects of NSSI and depression, highlighting the importance of considering the influence of comorbid mental disorders when investigating the cognitive underpinnings of such behaviors, especially in the context of prevalent cross-diagnostic phenomena like NSSI behaviors.

## Introduction

Non-suicidal self-injury (NSSI), prevalent in adolescents (1) and psychiatric populations (2,3), particularly in individuals with depression (4,5), involves deliberate self-harm behaviors (e.g., cutting, burning, scratching, and banging or hitting) without suicidal intent. NSSI’s lifetime prevalence is 17.2% among adolescents (6) and escalates to 40-80% among adolescents in clinical settings (7). Though distinct from suicidal behavior, NSSI is crucial to understand due to its strong association with future suicide risk (8–10). NSSI has been described as a maladaptive form of self-help (11), often used as a coping mechanism for intense emotional distress (12–15). Similar to suicidal actions, choosing to self-harm instead of less harmful coping solutions (e.g., talking to a friend, practicing mindfulness, or seeking professional counseling) reflects maladaptive decision-making (16). Our work, therefore, focuses on the decision characteristics of individuals with NSSI, aiming to identify the potentially multi-faceted cognitive dysfunctions behind NSSI decisions.

One early and influential hypothesis about NSSI is impulsivity (17), according to which impulsive individuals are more likely to engage in NSSI, due to attraction to the immediate emotional relief of self-harm and care less about its long-term negative consequences. Impulsivity is often measured by self-report scales such as the Barratt Impulsiveness Scale (18) and the UPPS Impulsivity Behavior scale (19), where it refers to personal traits including lack of forethought and carelessness. Consistent with the impulsivity hypothesis, self-reported impulsivity measures are often (though not always) higher in individuals with NSSI (17) and may predict the new onset of NSSI (20). However, the impulsivity hypothesis receives support rarely from behavioral measures of decision-making tasks (17,21), except when “impulsivity” is used as an umbrella term covering risky decision-making (21,22). In risky decision-making (23), someone who turns down an investment offer of 90% probability winning $1000 and 10% probability losing $2000 is not necessarily due to lack of forethought or carelessness, but due to aversion to risk (i.e., uncertainty in payoff) or loss.

What has emerged from the literature is that individuals with NSSI seems to make worse, less adaptive decisions than those without NSSI, in risky decision tasks ranging from Iowa Gambling Task (21), Cambridge Gambling Task (24), to Balloon Analogue Risk Task (BART) (22). This motivates us to consider NSSI behaviors as the result of risky decision-making (23), through which perspective individuals with NSSI would evaluate various emotion-coping options, each with its own set of risks, rewards, and losses. These individuals effectively outweigh the potential “benefits” (e.g., emotional relief) from self-harm against its physical and psychological costs (e.g., pain and shame) (25), and prefer it over less harmful solutions that nevertheless come with uncertain levels of risk, such as the challenges of potential stigma and accessibility issues (26,27). However, previous studies testing risky decision-making have often yielded inconsistent results in differentiating between individuals with and without NSSI (17,22,24,28). One reason for these mixed findings could be the inherent stochasticity of risky decision tasks, where different choices lead to variable experiences and thus exert different influences on subsequent decision behaviors, rendering simple behavioral measures less effective in capturing the complexity of individuals’ decision characteristics. Furthermore, the frequent comorbidity of NSSI with conditions like depression (29–31) adds another layer of complexity, potentially confounding the interpretation of the findings. Research on depression (often overlooking the comorbidity of NSSI) also has mixed findings: depressive moods decrease individuals’ risk-aversion (32), but patients with depression may exhibit different risk preferences in different risky decision tasks, such as lower risk-aversion in Iowa Gambling Task but higher risk-aversion in BART (33). These limitations highlight the necessity of using more advanced methods to understand NSSI-related decision characteristics.

In the present study, we investigated individuals with major depressive disorder (MDD) exhibiting NSSI behavior at least one episode within the past year (D+NSSI), contrasting them with MDD individuals without NSSI (D) and healthy controls (HC). Our approach involved the use of the BART (34) alongside computational modeling to provide a comprehensive understanding of NSSI-related decision characteristics and their modulation by depression. The BART simulates real-world risk-taking behavior by requiring participants to pump a virtual balloon for potential rewards (33). It not only assesses attitudes toward risk and loss, as widely known, but also delves into broader cognitive dimensions, such as prior beliefs about risk, learning from outcomes, and behavioral consistency under risk and uncertainty. In contrast, neither the Cambridge Gambling Task nor the Iowa Gambling Task assesses prior beliefs about risk. Additionally, the Cambridge Gambling Task cannot probe learning of risk from experience, as the risk is explicitly stated as part of the decision problem. Using computational modeling, we were able to disentangle these distinct cognitive dimensions from participants’ decision behaviors, by estimating the model parameters for each group based on the hierarchical Bayesian method. Furthermore, we developed a novel statistical method to isolate the effects of NSSI, depression level, and their interaction on model parameters. This allowed us to unravel the complex interplay of NSSI and depression in shaping individual decision characteristics, offering insights otherwise unattainable.

The existing literature does not allow us to fully expect the results of our study. One previous study tested BART on adolescents with or without NSSI history (22) found that the former made fewer pumps, which they interpreted as risk-aversion. We hypothesize patients with MDD who engage in NSSI behaviors are likely to display comparable patterns of risk-aversion and expect that depressive symptoms will interplay with the NSSI behaviors in the risk-related cognitive components, such as risk-aversion, loss aversion, and prior belief of risk.

## Methods and Materials

### Participants

From April 2022 to Sept 2023, we recruited 101 Chinese-speaking patients with depression from Peking University Sixth Hospital, including 48 outpatients and 53 inpatients. This cohort included 58 cases (43 inpatients and 15 outpatients) who engaged in NSSI behaviors at least one episode in the past 12 months (D+NSSI) and 43 cases (10 inpatients and 33 outpatients) without NSSI behaviors (D). Additionally, 157 healthy controls (HC) were recruited from two universities. All participants completed a custom questionnaire designed to collect detailed self-reported information on the presence and frequency of NSSI behaviors. All D+NSSI participants had engaged in NSSI behaviors for at least one episode in the past 12 months, whereas all D or HC participants had no history of self-injury in their lifetime. After quality control, 241 participants were included in the final data analysis. See Supplement for additional criteria for patient inclusion and exclusion and quality control details. The study was approved by the medical ethics committee of Peking University Sixth Hospital (Ethics Review No. 39 of 2021) and all participants provided informed consent.

Following previous studies (35), we used the term NSSI in more generally, referring to individuals who engaged in NSSI behaviors in the past year instead of more specifically to individuals with a separate diagnosis of NSSI disorder in the Diagnostic and Statistical Manual for Mental Disorders, Fifth Edition (DSM-5) (36). Nevertheless, supplementary analyses were performed to compare the subsets of participants who engaged in NSSI behaviors for at least five episodes and those fewer than five episodes in the past 12 months (*N=*30 for D+NSSI^≥5^ and *N=*27 D+NSSI^<5^).

### Psychiatric Assessment

The Zung self-rating depression scale (SDS) (37) and Zung self-rating anxiety scale (SAS) (38) were used to assess depression and anxiety levels, and raw scores were used. The SDS and SAS are both 20-item scales. Raw scores of 40 and 36 are typically considered indicative of clinical depression and anxiety, respectively (39,40). All assessments used Chinese versions of the scales, which were translated and validated for reliability and validity (41).

### Task and Design

On each trial of BART (Figure 1A), participants saw a virtual balloon on the screen and decided whether to pump (potentially increasing reward points) or collect points and end the trial. Each pump could increase the balloon’s size and thus the reward points by 1 but with the risk of popping and nullifying the points. Each trial thus presented a continual gamble between potential reward and loss. Each participant completed 60 trials of 30 red and 30 blue balloons, initially not knowing which color represented a higher risk. The higher-risk color would pop after 1 to 32 pumps (“32-balloon”) while the lower-risk color after 1 to 128 pumps (“128-balloon”). The number of pumps that maximizes expected reward is thus respectively 16 and 64 for the 32- and 128-balloons (Figure 1A, inset). Including these two different levels of risk in the experimental design allowed us to infer participants’ prior beliefs and learning as well as their attitudes towards loss and risk from their pumping decisions. See Supplement for more details.

**Figure 1.**
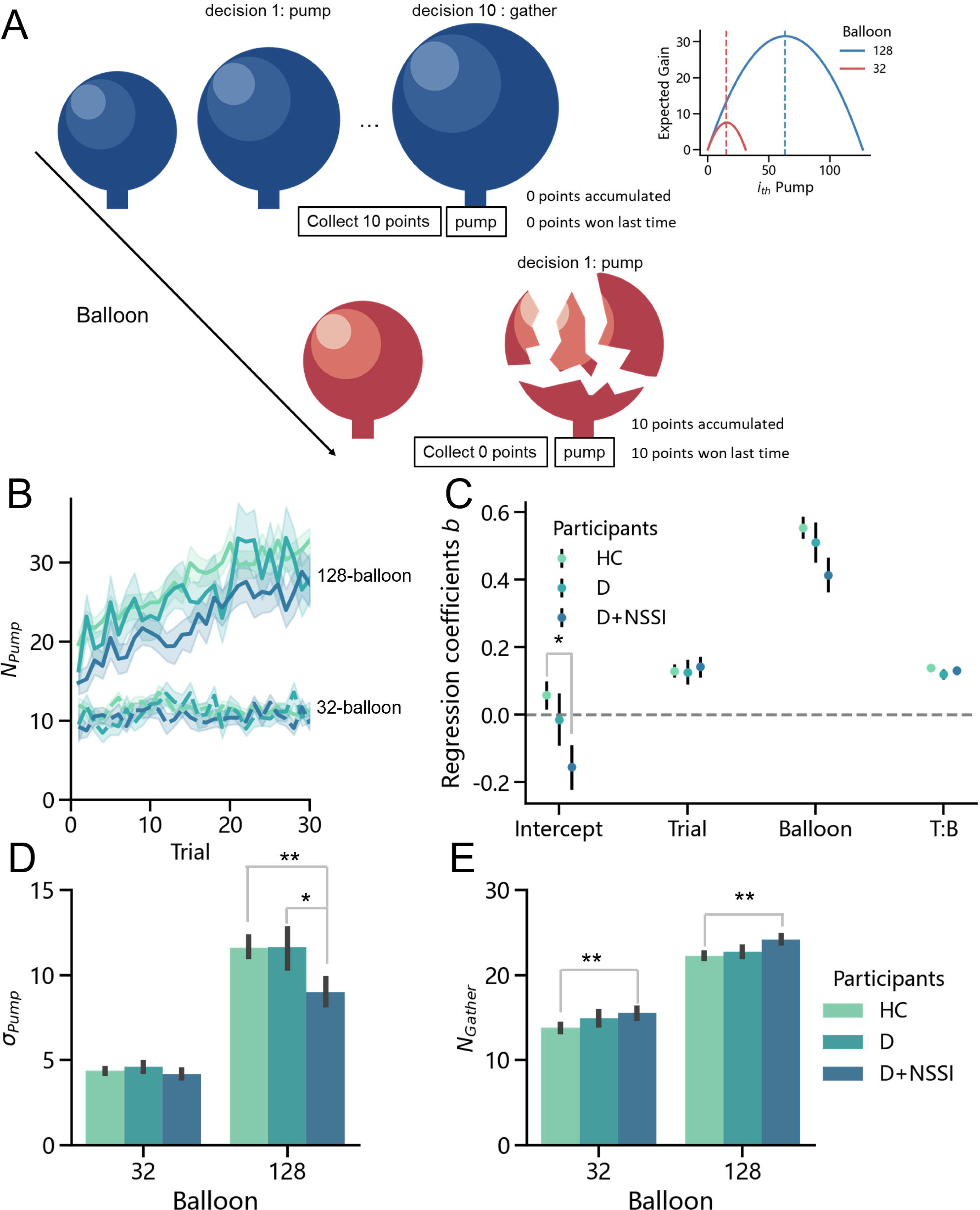
Balloon Analogue Risk Task (BART) and behavioral measures for the three groups of participants. **A**. Schematic representation of the BART, where at each moment participants choose to either pump the balloon to potentially accumulate one more reward point or to collect their accumulated points. If the balloon pops, the reward points accumulated for the balloon would be lost. The risk of balloon popping increases with each pump and varies with the color of the balloon. A new balloon starts after the old balloon popping or being collected. Each participant completed 30 red and 30 blue balloons (trials) in a random order. **Inset at top right**: Expected gain as a function of the number of pumps. The optimal numbers for the two balloon types (max pump before popping: 32 and 128), as indicated by the dashed vertical lines, are respectively 16 and 64. **B–E** plot the behavioral measures for the three groups of participants (*N=*241). Color codes: light-green for the 142 healthy controls (HC), blue-green for the 42 participants with major depressive disorder (MDD) but no NSSI behavior (D), and dark-blue for the 57 participants with MDD and NSSI (D+NSSI). **B**. How the mean number of pumps in non-popping trials (*N_pump_*) changed over trials. Curves in dashed lines and solid lines are respectively for the balloon types 32 and 128. Shadings denote 1 SE. **C**. Results of the linear mixed-effects model analysis on *N_pump_* predicted by group (intercept), trial, balloon, and the interaction of trial by balloon (T:B). The standardized coefficient *b* is plotted separately for each group (color coded). Error bars represent 1 SE of the coefficients. **D**. Standard deviation of *N_pump_* across trials (*σ_pump_*) for each balloon type and participant group. **E**. The number of non-popping (i.e., reward gathered) trials (*N_gather_*) for each balloon type and participant group. Error bars denote 1 SE. *: *p*<0.05. **: *p*<0.01. ***: *p*<0.001. The asterisks in **E** indicate a main effect between the HC and D+NSSI groups, regardless of the balloon type.

### Statistical Analysis

We conducted the following linear mixed-effects model (LMM) analysis on the number of pumps *N_pump_* in the non-popping trials to test the potential influence of major depressive disorder (D) and NSSI behavior (NSSI) on participants’ decision:

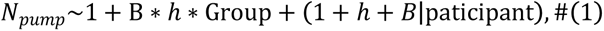

where B denotes the balloon type (–1 for “32-balloon” and 1 for “128-balloon”), *h* is the index of the balloon in the experiment, Group denotes categorical variables for the three participant groups, HC, D, and D+NSSI. The (|paticipant) denotes the random-effects terms that vary with participants. To further examine the potential different effects of NSSI<5 versus NSSI≥5, we conducted another LMM analysis:

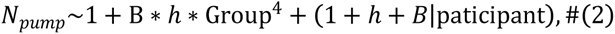

where Group^4^ denotes 4 participant groups, HC, D, D+NSSI^≥5^, and D+NSSI^<5^. All data were normalized on the group level before entering LMMs. All LMMs were implemented in R 4.0.3.

### Computational Modeling

#### The EMWV-VS model

The computational model we used to fit participants’ decision behaviors in BART (illustrated in Figure 2A), the EWMV-VS model, is an adaptation of the Exponential-Weight Mean–Variance (EWMV) model of Park et al. (42) to improve the reliability of parameter estimation (see Supplement for the model definition and its comparison with alternative models).

**Figure 2.**
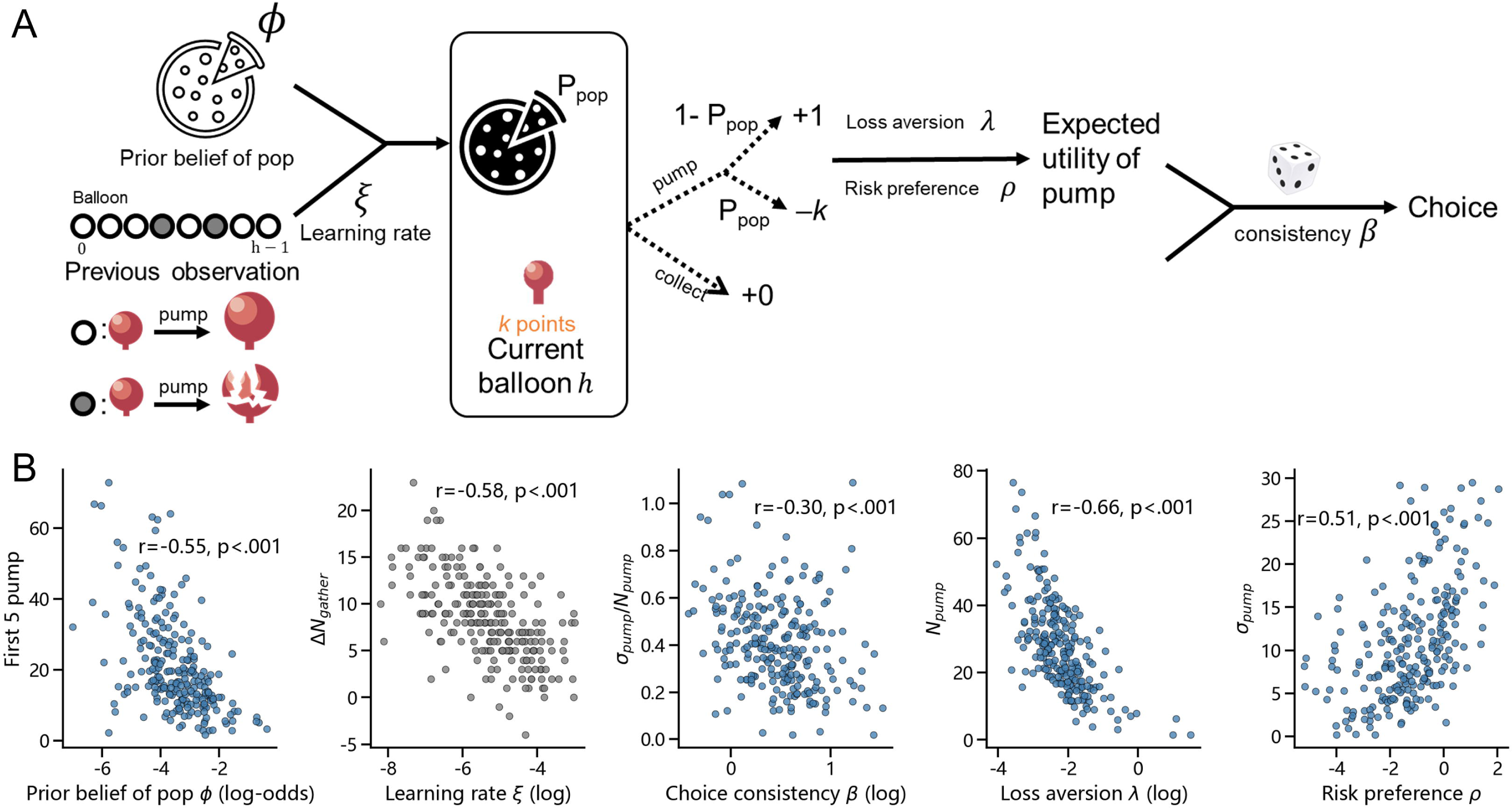
The computational model and its parameters’ behavioral implications. **A**. Illustration of the computational model used to simulate participant decisions in BART. The model agent integrates its prior belief of balloon popping (*φ*) with previous observations up to the last trial (i.e., balloon *h* − 1), resulting in the updated belief of *P_pop_*. How fast the prior belief is wiped off by increasing observations is determined by the learning rate (*ξ*). For the current balloon *h* that has accumulated *k* points, the agent is now faced with the decision of either pumping to gain 1 more point but with a chance of *P_pop_* losing all *k* points, or collecting the points. The agent’s loss aversion (*λ*) and risk preference (*ρ*) contribute to its expected utility of pumping the balloon. The probability of choosing pumping is further modulated by the agent’s choice consistency (*β*). **B**. Scatter plots showing how individuals’ model parameters (prior belief of pop *φ*, learning rate *ξ*, choice consistency *β*, loss aversion *λ*, and risk preference *ρ*) correlated with their behavioral measures in the BART. “First 5 pump” refers to the mean number of pump in the first 5 trials of the 128-balloons, which reflects the individual’s prior belief of pop. “Δ*N_gather_*” refers to the difference in the number of reward-gathered trials (*N_gather_*) between the 128- and 32-balloons, which reflects how much the individual learns from experience. The *N_pump_* and *σ_pump_*, as in Figure 1, respectively refer to the mean and standard deviation of the number of pumps, which and the combination of which are closely related to loss aversion, risk preference, and choice consistency. Data points are color-coded to indicate whether the behavioral measure is elicited from the 128-balloons (in gray-blue) or the difference between the 128- and 32-balloons (in gray). Each dot denotes one individual participant. The correlation coefficient (*r*) and p-value on each panel is based on Pearson’s correlation.

#### Effects-on-Parameters Analysis

To assess the joint effects of NSSI and depression severity on individuals’ decision characteristics, we developed an Effects-on-Parameters Analysis (EPA). Specifically, we built a hierarchical Bayesian model based on EWMV-VS, where each of the five parameters of EWMV-VS is generated from a linear combination of NSSI (as a binary variable of with or without NSSI behaviors), SDS, and their interaction term. The 15 coefficients were then estimated from participants’ choice data via hierarchical Bayesian modeling. For control analyses, we added age and education year as regressors to the group-level linear model. SDS, age and education year were normalized before entering the model.

## Results

### Demographic and clinical characteristics

A total of 241 participants were included in data analysis, including 142 healthy controls (HC), and 99 patients diagnosed with major depressive disorder (MDD), among whom were 57 cases with comorbid NSSI (D+NSSI) who engaged in NSSI behaviors at least one episode in the past 12 months and 42 reporting no history of NSSI in their lifetime (D). Participants’ gender, age, education year, self-rating anxiety and depression scales were recorded (Table 1). Because anxiety is highly comorbid with depression (43), there was a high correlation between the SDS and SAS scores in our participants with MDD (Pearson’s *r*=0.893, *p*<0.001), which made their effects practically undistinguishable. Therefore, we will report the effects of SDS only. Group Differences in Behavioral Measures

**Table 1.** Demographic and clinical characteristics.

| | Total<br>(N=241) | HC<br>(N=142) | D<br>(N=42) | D+NSSI<br>(N=57) | $\chi^2/F$ | p-value | Effect<br>size |
| --- | --- | --- | --- | --- | --- | --- | --- |
| **N by**<br>**gender** | | | | | $\chi^2(2) = 3.306$ | 0.191 | $V=0.017$ |
| Male | 78 (32.4%) | 49 (34.5%) | 16<br>(38.1%) | 13<br>(22.8%) |  |  |  |
| Female | 163 (67.6%) | 93 (65.5%) | 26<br>(61.9%) | 44<br>(77.2%) |  |  |  |
| **Age**<br>**(year)** | 19.80±2.36 | 19.30±2.05 | 21.10±2.<br>57 | 20.09±2.<br>54 | $F(2, 238)=10.803$ | <0.001 | $\eta^2=0.083$ |
| **Educatio**<br>**n year** | 13.49±2.11 | 13.11±1.77 | 14.43±2.<br>49 | 13.75±2.<br>36 | $F(2, 238)=7.313$ | <0.001 | $\eta^2=0.058$ |
| **SAS** | 39.66±14.69 | 31.33±6.30 | 45.90±12<br>.46 | 55.82±15<br>.48 | $F(2, 238)=123.718$ | <0.001 | $\eta^2=0.510$ |
| **SDS** | 45.34±16.83 | 34.93±8.00 | 51.90±11<br>.56 | 66.42±13<br>.96 | $F(2, 238)=198.908$ | <0.001 | $\eta^2=0.626$ |

*Note.* Demographic or scale scores are in mean ± standard deviation. SAS: self-rating anxiety scale; SDS: self-rating depression scale. $V$ : Cramér's $V$ , effect size for chi-squared test; $\eta^2$ : effect size for ANOVA.

### Group differences in *N_pump_*

Following the common practice of BART (34), a primary behavioral measure we used is participants’ number of pumps in non-popping trials (denoted *N_pump_*). Consistent with classical findings, for both levels of risk, participants pumped much less than the optimal number (Figure 1BC). Meanwhile, when we plot how the mean *N_pump_* of each participant group changed over trials (Figure 1B), we can see differences between different groups as well as trends of learning, at least for the 128-balloons. A linear mixed-effect model (LMM) analysis was performed on *N_pump_* to identify the effects of participant group (Intercept in Figure 1C), balloon type, and trial number as well as their interactions. This LMM analysis showed a significant main effect of participant group (*F*(2, 235.63)=3.68, *p*=0.027). Post-hoc analysis (multi-comparison corrected) further indicated that the D+NSSI group pumped fewer than the HC group (*b*=–0.213, *SE*=0.078, *p_holm_*=0.020, 95% CI [–0.401, –0.025]). There were no significant interactions between participant group with balloon type or trial number.

### Learning effects

We also observed significant learning effects for all groups. The mean *N_pump_* for 128-balloons was larger than that of 32-balloons (*F*(1, 232.18)=296.01, *p*<0.001). As the experiment proceeded, participants’ *N_pump_* increased overall (*F*(1, 231.74)=57.98, *p*<0.001), and changed differently under different balloon type (T:B in Figure 1C, interaction of balloon type by trial number, *F*(1, 8466.29)=340.76, *p*<0.001), with the *N_pump_* increasing over trials for 128-balloons (simple main effect, *b*=0.261, *SE*=0.018, *p*<0.001, 95% CI [0.225,0.296]) but staying constant for 32-balloons. We did not find significant differences between different group of participants in the learning effects (interaction of participants group by trial number, *F*(2, 231.84)=0.07, *p*=0.94).

### Group differences in *σ_pump_* and *N_gather_*

To provide a more comprehensive behavioral measurement for participants’ decision characteristics in BART, we also calculated the standard deviation of *N_pump_* across trials (denoted *σ_pump_*) and the number of non-popping (i.e., reward gathered) trials (denoted *N_gather_*) for each balloon type and participant group (Figure 1DE). Mixed-design ANOVAs for these two measures showed a significant interaction of participant group by balloon type for *σ_pump_* (*F*(2, 238)=3.10, *p*=0.047, 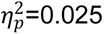), whereas no significant interaction was observed for *N_gather_* (*F*(2, 238)=0.476, *p*=0.622, 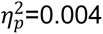). We also observed significant main effects of participant group for *σ_pump_* (*F*(2, 238)=3.39, *p*=0.035, 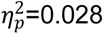) and *N_gather_* (*F*(2, 238)=4.88, *p*=0.008, 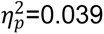), other than the obvious differences between different balloon types (*F*(1, 238)=187.33, *p*<0.001, 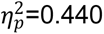 for and *F*(1, 238)=679.75, *p*<0.001, 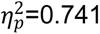 for *N_gather_*). According to simple main effects, participants in different groups had significantly different *σ_pump_* for 128-balloons (*F*(2, 238)=3.44, *p*=0.034), but not for 32-balloons (*F*(2, 238)=0.73, *p*=0.484; Figure 1D). *Post-hoc* comparisons further indicates that for 128-balloons, participants in the D+NSSI group had smaller *σ_pump_* than both the HC (difference *M*=–2.634, *SE*=0.761, *t*=–3.46, *p_holm_*=0.004) and D groups (difference *M*=2.647, *SE*=0.987, *t*=–2.68, *p_holm_*=0.038). Compared to the HC group, participants in the D+NSSI group also had larger σ_*pump*_ (difference *M*=1.844, *SE*=0.597, *t*=3.09, *p_holm_*=0.007; Figure 1E).

Together, these behavioral measures of BART suggest that the HC, D, and D+NSSI groups had distinct decision patterns, though all groups similarly learned from their experiences. Findings were comparable when age or education year was controlled for in these models (see Supplemental Statistical Analysis). Because all behavioral measures are the joint consequence of multiple cognitive components (e.g., learning, risk attitude, etc.), the underlying cognitive differences remained elusive. We will disentangle the cognitive dimensions below, with the help of computational modeling.

### Five Cognitive Dimensions Decomposed by the Computational Model

The computational model we used to fit participants’ decision behaviors is an adaptation of the Exponential-Weight Mean–Variance (EWMV) model of Park et al. (42), termed EWMV-VS (see Methods), which outperforms its precedent in predictive power (see Supplemental Table S1) and has a good parameter recovery performance (Supplemental Figure S1).

When fitted to participants’ decision behaviors, the five parameters of the EWMV-VS model—prior belief of pop *φ*, learning rate *ξ*, choice consistency *β*, loss aversion *λ*, and risk preference *ρ*—can thus represent participants’ characteristics on five distinct cognitive dimensions (see Figure 2A and its legend for details). Some of these model parameters have intuitive behavioral implications, as indicated by the correlations between individual participants’ model parameters and their behavioral measures (Figure 2B).

### Revealing Group-Level Cognitive Differences: Hierarchical Bayesian Modeling

To assess differences in each cognitive dimension across the three groups, we fit the EWMV-VS model to participants’ decision behaviors using hierarchical Bayesian modeling (see Supplement). Among the five parameters (Figure 3A), we observed significant group differences in the parameters of choice consistency and risk preference. In particular, the choice consistency decreased in the depressed-only patients (i.e., the D group), both the D+NSSI (difference *M*=0.24, 95% HDI [0.03,0.44]) and HC (difference *M*=0.19, 95% HDI [0.02,0.36]) groups exhibited a higher choice consistency. The D+NSSI group also had a lower risk preference than the HC group (difference *M*=–0.75, 95% HDI [–1.5, –0.029]), indicating a more cautious approach to risk-taking.

**Figure 3.**
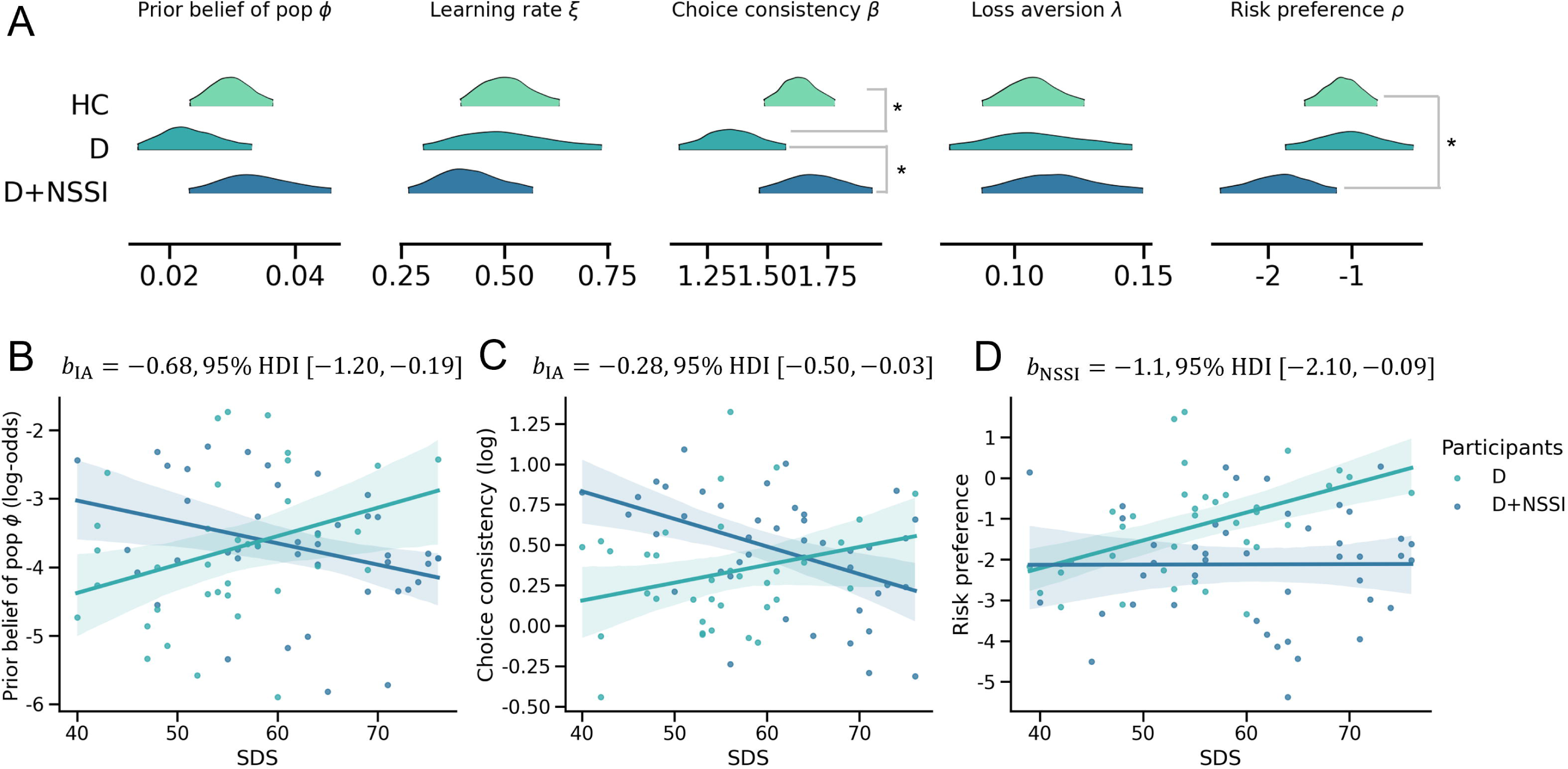
Estimated model parameters for different participant groups. **A**. The 95% highest density interval (HDI) of the posterior density of the group-level model parameters estimated from hierarchical Bayesian model fitting. Significant differences between groups were found for choice consistency (*β*) and risk preference (*ρ*). See text. **B–D** show the joint effects of NSSI and depression severity (as measured by the Self-Rating Depression Scale, SDS) on model parameters, where the participants in analysis were limited to the D and D+NSSI groups with an overlapping range of SDS. Each dot denotes one individual participant. The shading on the regression line denotes the 95% HDI. **B**. Interaction (*b_IA_*) between NSSI and SDS on prior belief of pop (*φ*). Higher depression severity was associated with a lower belief in pop likelihood for the NSSI group, while the reverse for the non-NSSI group. **C**. Interaction (*b_IA_*) between NSSI and SDS on choice consistency (*β*). Higher depression severity was associated with less consistent choices in the NSSI group, while the reverse for the non-NSSI group. **D**. Main effect of NSSI (*b_N_*) for risk preference (*ρ*). The NSSI group was less tolerant of risk than the non-NSSI group. The asterisk (*) denotes that the 95% HDI of the group difference did not contain 0 (similar to 0.05-level significant differences in the frequentist statistics such as *t* tests).

These findings, rooted in our model analysis, echo the behavioral patterns described earlier, specifically the reduced variability in the number of pumps (*σ_pump_*) among the D+NSSI group (Figure 1D). While behavioral data alone might suggest this decreased variation could stem from diminished learning, implying more uniform behavior from early to late trials, our model-based analysis mostly rules out this possibility.

### Unraveling NSSI and Depression Interplay: Effects-on-Parameters Analysis

We noted that the D+NSSI group had a significantly higher depression severity than the D group (as indicated by the SDS score in Table 1), despite both groups being diagnosed with MDD and the majority had SDS scores above the cut-off point for clinical significance (44). To distinguish the impact of NSSI from depression severity, we combined two strategies. First, we selected from the D and D+NSSI groups the participants whose SDS fell into an overlapping range (SDS range [39, 76], with 37 D and 44 D+NSSI, see Table 2). Second, we developed a novel Effects-on-Parameters Analysis (EPA, see Methods), which includes NSSI, SDS, and their interaction as linear predictors of the EWMV-VS model’s parameters in hierarchical Bayesian modeling, providing a granular view of their joint effects on individuals’ decision characteristics.

**Table 2.**
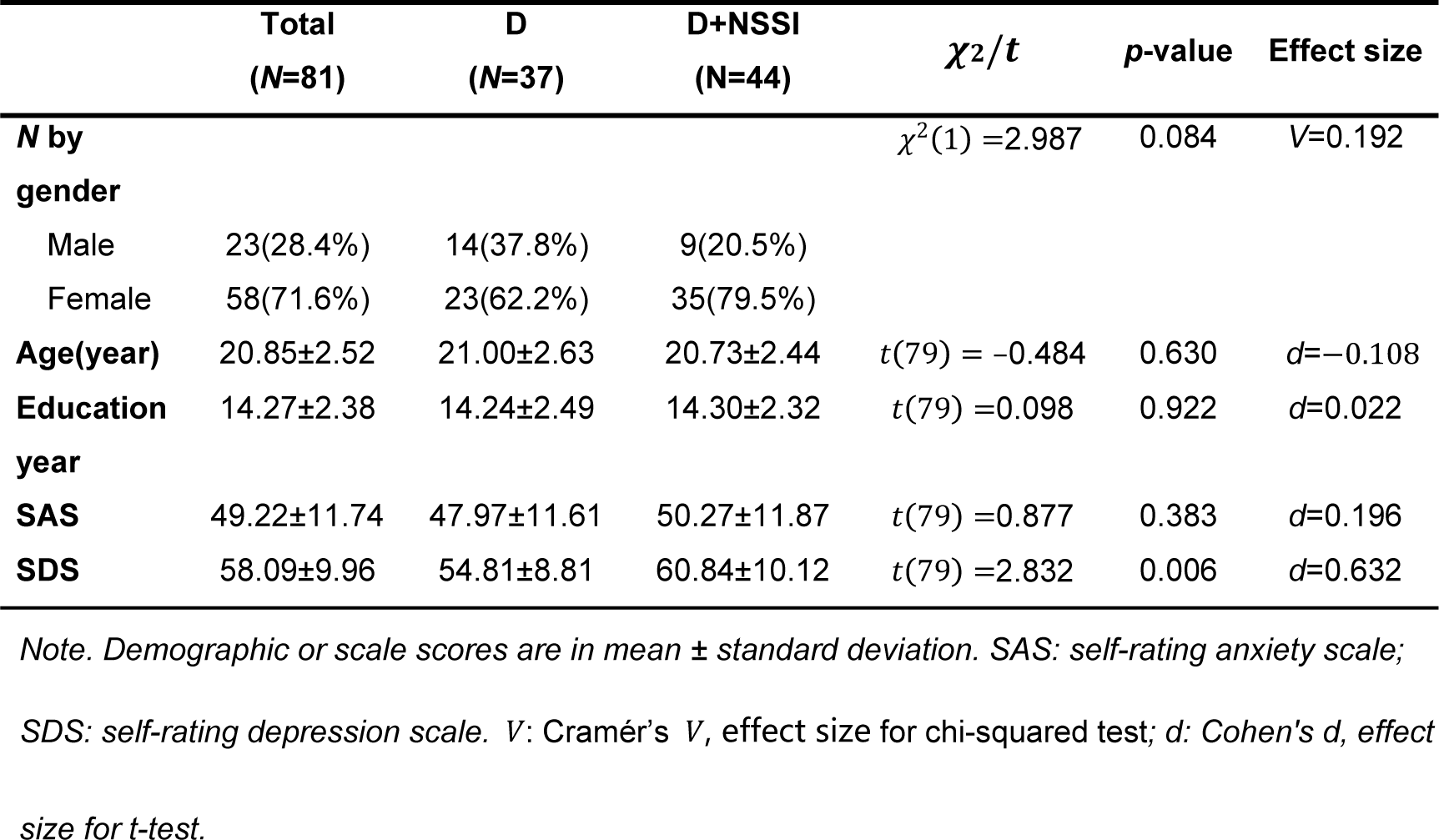
Participants included in the effects-on-parameters analysis.

By applying EPA to the 81 participants’ decision data, we found that NSSI was still associated with a decrease in risk preference (*M*=–1.1, 95% HDI [–2.1, –0.09]), even when depression severity was controlled (Figure 3D). SDS had no significant main effect on risk preference, nor did its interaction with NSSI. NSSI and SDS jointly influenced the prior belief of pop and choice consistency in a more complicated way, showing as interaction effects. Specifically, the prior belief of pop increased with SDS for the depressed-only participants but decreased with SDS for those with NSSI behaviors (Figure 3B; *M*=–0.68, 95% HDI [–1.2, –0.19]). Choice consistency rose with SDS among the depressed-only participants but fell among those with NSSI (Figure 3C; *M*=–0.28, 95% HDI [–0.5, –0.03]). These effects were robust, which were similar when the effects of age and education year were controlled (Supplemental Figures S2 and S3).

In sum, participants with NSSI behaviors were more risk-averse. However, unlike those with depression only, as their depressive symptoms intensified, these participants demonstrated a decreased prior belief of pop and lower choice consistency, implying a perception of the environment as less risky and a tendency towards more random choices.

### Similar Decision Behaviors in MDD Patients with NSSI<5 and NSSI≥5

The D+NSSI group in our analyses above comprised cases with varying frequencies of NSSI. To test the possibility that they might have different decision characteristics, we performed supplementary analyses where the D+NSSI group was divided into two sub-groups according to the self-reported NSSI frequency in the past 12 months, denoted D+NSSI^<5^ and D+NSSI^≥5^ (27 and 30 participants, respectively).

In Figure 4A, we visualize how the mean number of pumps in non-popping trials (*N_pump_*) changed over trials for the four participant groups. To examine the effects of NSSI<5 versus NSSI≥5, we first performed an LMM analysis on *N_pump_* similarly as before, except that the regressor of participant group included four instead of three categories (see Methods). All the significant effects identified in the previous LMM were replicated in this new LMM (Figure 4B, see Supplement for details). Hierarchical Bayesian modeling analysis for four participant groups also suggests that the NSSI≥5 and NSSI<5 groups had similar model parameters (Figure 4C and Supplemental Table S2). A further modeling analysis that compared a 3-group model with a 4-group model (see Supplement and Supplemental Table S3) also provides support for this lack of differences.

**Figure 4.**
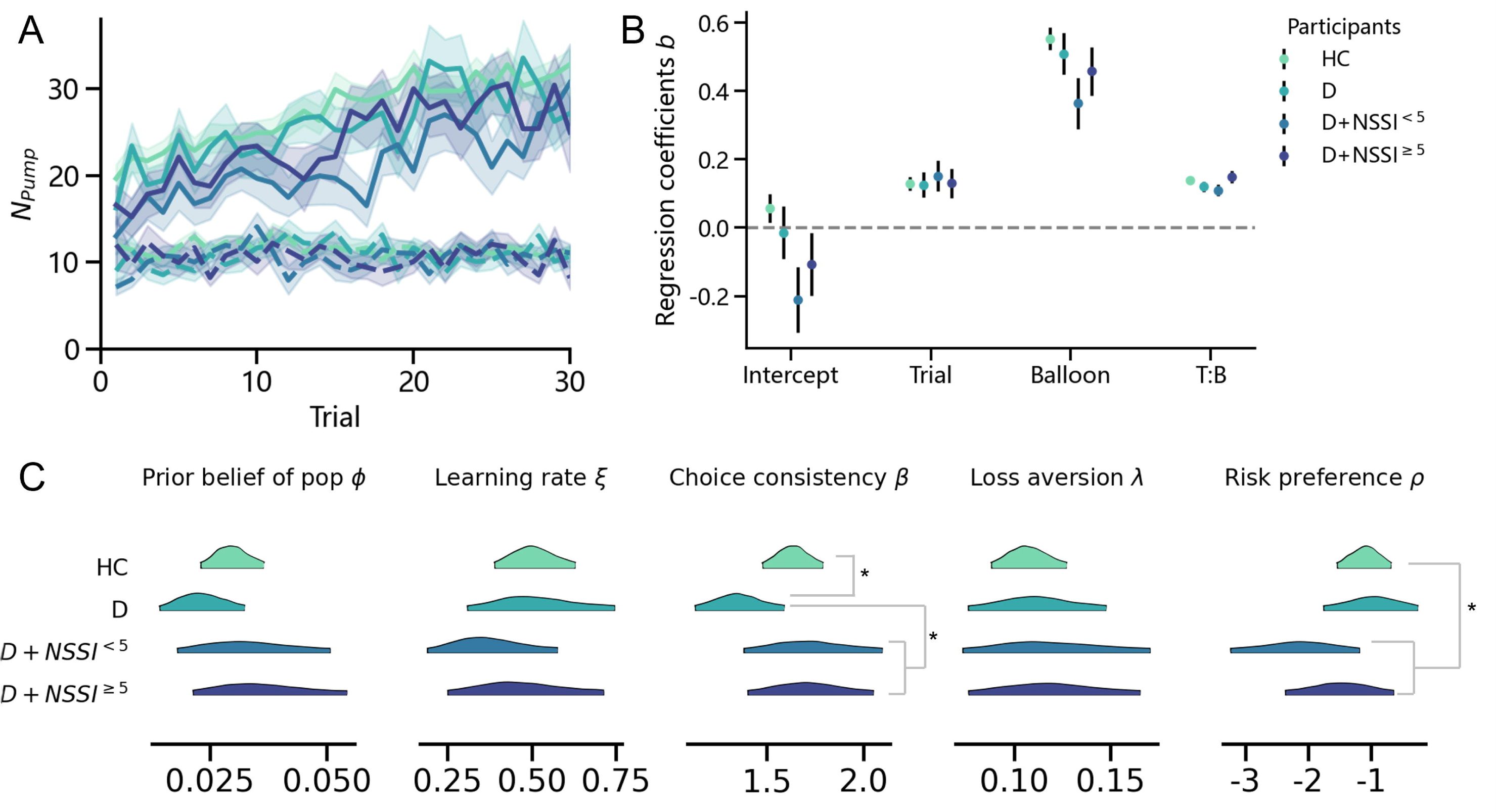
Behavioral measures and estimated model parameters for four participant groups. **A**. How the mean number of pumps in non-popping trials (*N_pump_*) changed over trials for four participant groups. Curves in dashed lines and solid lines are respectively for the balloon types 32 and 128. Shadings denote 1 SE. **B**. LMM model results. b is the standardized coefficient of the linear mixed-effects model. Error bars represent 1 SE of the coefficients. **C**. The 95% highest density interval (HDI) of the posterior density of the group-level model parameters estimated from hierarchical Bayesian model fitting. The asterisk (*) denotes that the 95% HDI of the group difference did not contain 0 (similar to 0.05-level significant differences in the frequentist statistics such as *t* tests). No significant differences were found for the parameters of D+NSSI^<5^ and D+NSSI^≥5^ groups. See text. *: *p*<0.05. **: *p*<0.01. ***: *p*<0.001. Asterisks on the left or right side of an ensemble of coefficients in **B** denote significant deviations from 0 for all these coefficients.

## Discussion

Our finding that individuals with NSSI behaviors are more risk-averse instead of more “impulsive” as commonly perceived is consistent with that of Dillahunt et al. (22), who found adolescents with self-injury history made fewer pumps in BART. Despite seemingly counterintuitive, there has been other evidence that NSSI behaviors may be associated with risk-aversion (45,46). Such risk-aversion can be understood from the perspective how NSSI serves as an emotional regulation mechanism (47) . Individuals engaging in NSSI often seek relief from intense negative emotions or aim to achieve a positive emotional state, suggesting a controlled (i.e., less risky), albeit maladaptive, response to distress. In contrast, pumping the balloons in the BART produces less controlled (i.e., more risky) outcomes. This mirrors their preference for self-injury, a harmful yet certain means of emotional relief, over less harmful coping strategies with uncertain (“risky”) outcomes. Our analysis further reveals reduced exploratory behavior in individuals with NSSI, characterized by smaller variations in balloon pumps (σ_*pump*_) and higher choice consistency (β), without an impairment in learning rate (ξ). Both the aversion to risk and decreased exploration reflect limited adaptive coping strategies (12).

Contrary to previous reports of risk-aversion using BART in depression (33), we did not observe fewer balloon pumps among depression-only individuals. This discrepancy underscores the necessity of accounting for sample heterogeneity, given that NSSI was not examined in prior research on depression. The Effects-on-Parameters analysis we developed enables us to further disentangle the effects of NSSI and depression, revealing interaction effects in the prior belief of risk and choice consistency. Choice consistency is related to random exploration, which was reported to be lower in depressed patients (48). Such intricate interplay between NSSI and depression may have led to the controversial results documented in previous studies (17). Furthermore, given NSSI’s strong predictive value for future suicidal behavior (8,9), managing depressive symptoms in individuals with NSSI could be a key preventative measure.

Previous research suggests that risk taking differs between younger and older adults (49) and that adolescents are more likely to engage in risky behaviors than adults (50,51). The age range in our study was relatively narrow but covered mid-late adolescence to emerging adults. Considering the significant (though small) differences between groups in age, we included age and its highly correlated educational level as confounding variables in supplementary modeling analysis to remove their potential impact, where our major findings still hold. Besides, the D+NSSI group in our study, despite being younger than the D group, exhibits stronger risk-aversion, which is in the direction reverse to the age-related effects suggested by previous research. Therefore, the likelihood of our findings being age-driven is relatively low.

The following limitations may be addressed in future research. First, due to time constraints of outpatient visits, we did not diagnose borderline personality disorder (BPD), a condition in which NSSI is prevalent (3,30), and therefore did not exclude the potential BPD patients from our study. According to an epidemiologic survey in China, approximately 11.49% of 258 outpatient cases with MDD met the BPD diagnosis criteria (52). The estimated potential BPD cases in our patient cohort is thus approximately 5 to 6. Although this number is small, controlling for BPD comorbidity would be ideal for future research. Second, though no significant differences were observed between the NSSI≥5 and NSSI<5 groups, our findings may not necessarily apply to the persistence of NSSI behaviors that can lead to significant functional impairment. Expanding the study to include a larger sample (including those with NSSI behaviors but without mental disorder) and structured scales for assessing the frequency and severity of NSSI would enhance the robustness of the findings.

Additionally, the prevalence of NSSI behavior is generally higher among female adolescents than males (53,54). Females reported significantly greater psychological distress and lower levels of sensation seeking and positive urgency compared to males (54). While there is no conclusive link established between NSSI behaviors and gender, further exploration of demographic factors, such as gender and age, could provide deeper insights.

In conclusion, NSSI behaviors manifest a general aversion to risk. However, as depressive symptom severity increases, individuals perceive a decrease in environmental risks and demonstrate more unpredictable behaviors. This suggests a complex interaction between depressive symptoms and NSSI behaviors in decision-making. On one hand, overestimating risks in unfamiliar environments may diminish the likelihood of exploring alternative means to cope with emotions. On the other hand, depression amplifies engagement in risky behaviors.

## Acknowledgments

We extend our gratitude to all participants in this study. This study was supported by the Clinical Medicine Plus X – Young Scholars Project, Peking University the Fundamental Research Funds for the Central Universities (PKU2023LCXQ023) for HY and HZ. HY was partly supported by the National Natural Science Foundation of China (82071505). HZ was partly supported by the National Natural Science Foundation of China (32171095), and funding from Peking-Tsinghua Center for Life Sciences.

## Conflicts of interest

The authors have no conflicts of interest to declare.

## Notes

### Competing Interest Statement

The authors have declared no competing interest.

### Summary of Updates

In this revision, we have: 1.Corrected the inaccuracies regarding the DSM-5 frequency criterion for NSSI disorder. 2.Included the detailed explanation on the advantages of using the BART in our study. 3.Made all the necessary textual changes to enhance clarity and readability.

