## Supplemental Materials for "Cognitive Phenotype Shifts in Risk-Taking: Interplay of Non-Suicidal Self-Injury Behaviors and Intensified Depression"

### Supplemental methods and results

#### Additional inclusion and exclusion criteria for participants

Additional criteria for patient inclusion were: a) aged 16–25; b) diagnosed with MDD according to the International Classification of Diseases, Tenth Edition (ICD-10). Exclusion criteria included: a) engaging in suicidal self-injury; b) current diagnosis of other mental disorders such as pervasive developmental disorders, psychotic disorders, manic episodes, substance dependence, or obsessive-compulsive disorders; and c) presence of severe or unstable physical illness. Fourteen participants originally recruited as HC were excluded for a history of NSSI behavior. One HC and one D+NSSI participants were excluded for almost identical responses in each trial of the decision task. One D participant was excluded for missing data.

#### Task and design details

In the BART (Figure 1A), each trial presents participants with a virtual balloon on their computer screen. Participants can press a specific key to “pump” the balloon, which adds one point to their prize pool for each pump. However, every pump also carries a risk of causing the balloon to pop. If the balloon pops, the points in the prize pool for that balloon are lost, and the trial ends. Thus, participants must choose between pumping the balloon to potentially increase their reward point or collecting the accumulated points to conclude the trial safely. The total points collected across all trials would be converted into monetary compensation at the end of the experiment.

Each participant completed 30 trials with red balloons and 30 with blue balloons, presented in a random order. The two different colors represent two levels of risk of popping, corresponding to different optimal numbers of pumps to maximize expected reward (see Inset in Figure 1A). One color of balloon is associated with an equal chance of popping on any pump between the 1st and the 32nd (“32-balloon”), while the other between the 1st and 128th (“128-balloon”). Following this design, the number of pumps that maximizes expected reward is respectively 16 and 64 for the 32- and 128-balloons (Figure 1A, inset). In the illustration of Figure 1A, red balloons are 32-balloons and blue balloons are 128-balloons. The real color-risk correspondence was counterbalanced between participants. Participants were not informed the probability of popping or which balloon would be riskier, and therefore had to learn from their own experience in the experiment.

The experiment consisted of three blocks of 20 trials: the first block was a randomized mix of 10 red and 10 blue balloon trials, and the subsequent two blocks consisted of 20 trials of each balloon color. The order of the last two blocks was counterbalanced across participants. In total, it took approximately 10 minutes.

The presentation and data recording of the BART task were controlled by python, implemented via PsychoPy 2021.1.4.

#### Supplemental statistical analysis: controlling age and education year

The three group of participants had small but statistically significant differences in age and education year. To exclude the possibility that the behavioral differences we found between different groups was due to these demographic differences, we performed supplemental linear mixed-effects model (LMM) analyses, which are similar to the LMM analyses reported in the main text but include age or education year as the covariates.

$$\begin{aligned} N_{pump}\sim1+Age*B*h*Group+\left( 1+h+B | \mathrm{paticipant} \right),\#\left( S1 \right) \end{aligned}$$

$$\begin{aligned} N_{pump}\sim1+Age*B*h*\mathrm{Group}^{4}+\left( 1+h+B | \mathrm{paticipant} \right),\#\left( S2 \right) \end{aligned}$$

T$\begin{aligned} N_{pump}\sim1+Edu*B*h*Group+\left( 1+h+B | \mathrm{paticipant} \right),\#\left( S3 \right) \end{aligned}$

$$\begin{aligned} N_{pump}\sim1+Edu*B*h*\mathrm{Group}^{4}+\left( 1+h+B | \mathrm{paticipant} \right),\#\left( S4 \right) \end{aligned}$$

where Age is the normalized age variable, Edu is normalized education year. As in the main text, $B$ denotes the balloon type (–1 for “32-balloon” and 1 for “128-balloon”), $h$ is the index of the balloon in the experiment, Group denotes categorical variables for the three participant groups, HC, D, and D+NSSI, and $\mathrm{Group}^{4}$ denotes 4 participant groups, that is, HC, D, D+NSSI^≥5^, and D+NSSI^<5^. The $\left( \cdot| \mathrm{paticipant} \right)$ denotes the random-effects terms that vary with participants.

**Age-controlled LMM results**. Similar to the LMM results in the main text, for three-group LMM that controls age (Eq. S1), we found significant effects of participant groups (*F*(2, 232.20)= 4.09, *p* = 0.018), balloon type (*F*(1, 228.70)= 255.13, *p* < 0.001), trial number (*F*(1, 229.36)= 43.59, *p* < 0.001), as well as the interaction of balloon and trial number (*F*(1, 8477.68)= 294.93, *p* < 0.001). Age had no significant main or interaction effects on participants’ choice. The results of the four-group age-controlled LMM (Eq. S2) were also the same as those of the three-group LMM, with balloon type (*F*(1, 226.13)= 209.88, *p* < 0.001), trial number (*F*(1, 225.64)= 40.90, *p* < 0.001), and the interaction of balloon and trial number (*F*(1, 8463.30)= 258.07, *p* < 0.001), except that the main effect of participant groups was only marginally significant (*F*(3, 229.95)= 2.47, *p* = 0.063).

**Education-controlled LMM results**. Similar to the LMM results in the main text, for three-group LMM that controls education (Eq. S3), we found significant effects of participant groups (*F*(2, 232.92)= 3.61, *p* = 0.029), balloon type (*F*(1, 229.88)= 272.93, *p* < 0.001), trial number (*F*(1, 231.83)= 45.81, *p* < 0.001), as well as the interaction of balloon and trial number (*F*(1, 8481.65)= 321.81, *p* < 0.001). We also found a significant three-way interaction of education year, balloon and trial number (*F*(1, 8489.33)= 5.18, *p* = 0.023). Similar to the age-controlled analysis, in the four-group education-controlled LMM (Eq. S4), there were significant effects of balloon type (*F*(1, 226.80)= 220.25, *p* < 0.001), trial number (*F*(1, 227.00)= 44.19, *p* < 0.001), and the interaction of balloon and trial number (*F*(1, 8460.04)= 278.71, *p* < 0.001), though the main effect of participant groups failed to reach significance (F(3, 230.50)= 2.06, p = 0.106). As in the three-group analysis, there was a significant three-way interaction of education year, balloon and trial number (*F*(1, 8444.79)= 4.43, *p* = 0.035).

#### The definition of the EWMV-VS model

Below we first introduce the basic modeling idea and specific assumptions of EWMV, and then our adaptation—EWMV with variance standardization (EWMV-VS).

**Basic modeling idea**. After comparing a comprehensive set of computational models they constructed for BART, Wallsten et al. (1) found that human participants’ behaviors in BART are best captured by a model with three key assumptions for the belief updating and decision processes: 1) the agent believes that the probability of balloon popping could vary with balloon colors but is constant across different pumps for the same color of balloon, 2) the agent updates the probability of pop before each trial based on previous experience with popping and successful inflating, and 3) each agent has specific attitudes towards loss or risk. Following this stream of assumptions, Park et al. (2) further proposed the EWMV model, which has closely-related mathematical forms and a comparable predictive power, but whose parameters can be more reliably recovered from human behavior, thus allowing for a more accurate characterization of individual differences.

**EWMV**. According to the EWMV model, the agent starts with a specific prior probability (parameter $\phi$) for balloon popping, and updates its belief based on the numbers of popping and successful inflating previously observed for the balloons that matched the color of the current balloon. In the agent’s belief updating process, the relative weights for the prior probability and experience are approximated by exponential weighting functions:

$$\begin{aligned} p_{h}=\phi\exp\left( -\xi\sum_{h^{'}=1}^{h-1} m_{h^{'}} \right)+\left( 1-\exp\left( -\xi\sum_{h^{'}=1}^{h-1} m_{h^{'}} \right) \right)P_{h-1}.\#\left( S5 \right) \end{aligned}$$

Here $p_{h}$ denotes the pop probability per pump for the *h*-th balloon in its color, ξ is the parameter for learning rate. The higher the $\xi$, the faster the prior belief is washed out by experience. With $m_{\left( \cdot\right)}$ denoting the number of pumps for a specific balloon indicated by the subscript, $\sum_{h^{'}=1}^{h-1} m_{h^{'}}$ is the total number of pumps made to all previous $h-1$ balloons in the same color. For these $h-1$ balloons, $P_{h-1}$ is the observed pop probability per pump:

$$\begin{aligned} P_{h-1}=\frac{\sum_{h^{'}=1}^{h-1} d_{h^{'}}}{\sum_{h^{'}=1}^{h-1} m_{h^{'}}},\#\left( S6 \right) \end{aligned}$$

where $d_{h}$ indicates whether balloon $h$ finally popped (1) or not (0).

Suppose the agent has pumped the current balloon for $k$ times, accumulating $k$ points. At this moment, pumping corresponds to a gamble with probability ${1-p}_{h}$ to gain 1 more point but probability $p_{h}$ to lose the $k$ points; collecting is to win or lose nothing. To decide whether to pump or to collect the accumulated points, the agent calculated the expected utility of pumping relative to collecting:

$$\begin{aligned} U_{h,k}=\left[ \left( 1-p_{h} \right)-p_{h}\lambda k \right]+\rho p_{h}\left( 1-p_{h} \right)\left( 1+\lambda k \right)^{2}.\#\left( S7 \right) \end{aligned}$$

The first part of the equation, $\left[ \left( 1-p_{h} \right)-p_{h}\lambda k \right]$, is the sum of expected gain and expected subjective loss, with loss scaled by the loss aversion parameter $\lambda$, that is, how many units of gain one unit of loss equals to. With such subjective loss considered, the gamble is effectively a payoff distribution of probability ${1-p}_{h}$ to receive 1 point and probability $p_{h}$ to receive $-\lambda k$ points, whose variance is $p_{h}\left( 1-p_{h} \right)\left( 1+\lambda k \right)^{2}$ points. This variance, multiplied by the risk preference parameter $\rho$, constitutes the second part of the equation, the risk discounting term. A positive $\rho$ would imply preference for risk (i.e., more varied outcomes), while a negative $\rho$ would imply risk aversion.

The agent's probability of choosing pumping is a softmax function of this expected utility:

$$\begin{aligned} r_{h,k}=\frac{1}{1+e^{-\beta U_{h,k}}},\#\left( S8 \right) \end{aligned}$$

where the parameter $\beta$ characterizes choice consistency. The final choice of pumping or collecting is a Bernoulli random variable based on $r_{h,k}$.

**EWMV with variance standardization**. When the probability of balloon popping is relatively small (e.g., 1/32 and 1/128), there could be strong correlation between the mean and variance terms in the computation of expected utility (Eq. 5). As a result, the risk preference parameter $\rho$ cannot be accurately estimated. Similar issues would still occur if the risk preference parameter is incorporated in expected utility not as the scaling factor of variance but through the more classical form of a utility function (3,4). To alleviate this issue, we adopt the standardization method (4) to perform variance standardization on the expected utility:

$$\begin{aligned} U_{h,k}=\frac{\left[ \left( 1-p_{h} \right)-p_{h}\lambda k \right]+\rho p_{h}\left( 1-p_{h} \right)\left( 1+\lambda k \right)^{2}}{\sqrt{p_{h}\left( 1-p_{h} \right)}\left( 1+\lambda k \right)}.\#\left( S9 \right) \end{aligned}$$

The standardized $U_{h,k}$ is then entered into Eq. 6 to obtain the probability of choosing pumping. Similar to EWMV, there are five free parameters in our EWMV-VS model: prior belief of popping $\phi$, learning rate $\xi$, choice consistency $\beta$, loss aversion $\lambda$, and risk preference $\rho$.

**Model Fitting**. Hierarchical Bayesian modeling was used for parameter estimation, which allows more accurate estimation of group-level effects. The model parameters were estimated from participants’ choice data using the Markov Chain Monte Carlo method implemented by the PyMC package (version 5.9.2) on python 3.11. Four independent chains were run, each with 1000 samples after an initial burn-in of 1000 samples, which resulted in a good convergence of $\hat{R}\leq1.01$ for all estimated parameters. The 95% highest density intervals (HDI) were calculated for the group-level effects.

#### Testing the performance of the EWMV-VS model

As described in the Methods of the main text, we introduced variance standardization (4) into the Exponential-Weight Mean–Variance (EWMV) model of Park et al. (2) to improve the accuracy of parameter estimation. The resulting new model is called EWMV with variance standardization (EWMV-VS). Before using it to fit human participants’ behaviors, we performed additional analyses to ensure that 1) its fitting performance is at least as good as the original EWMV model and 2) its parameters can be accurately estimated from human behavioral data.

**Model comparison**. Model comparison results for a comprehensive set of computational models had been reported in previous research where the EWMV model was developed (1,2), according to which the EWMV model was the one best-fitting human behavioral data in BART. Here we compare our EWMV-VS model with EWMV model, and the winning model in Wallsten’s work (see Park et al., 2021; Wallsten et al., 2005 for further detail).

We fit Wallsten’s model, EWMV and EWMV-VS models separately to participants’ data using hierarchical Bayesian modeling, with all 241 participants as one group. The expected log pointwise predictive density (ELPD), assessed through the leave-one-out cross-validation (LOO) method, was used to estimate the predictive performance (i.e., goodness-of-fit corrected for over-fitting) of each model (5). Larger value of ELPD indicates better predictive performance. The model fitting and the computation of ELPD were implemented by the pymc (5.92) and arviz (0.16.1) packages on python 3.11.5. As shown in Table S1, the EWMV-VS model outperformed Wallsten et al.’s best model and the EWMV model in the predictive performance.

**Model recovery analysis**. We performed a model recovery analysis to evaluate how accurate the EWMV-VS model’s parameters can be estimated from behavioral data. In the model recovery analysis, we first used the EWMV-VS model with specific sets of parameters to generate simulated data and then fit these data to see how well the estimated parameters agreed with the true parameters. In particular, we calculated the mean and variance for the EWMV-VS parameters estimated for the real participants, separately for the three groups of participants, and passed them into Gaussian distributions to randomly generate 90 sets of model parameters, 30 for each group. The EWMV-VS model with a specific set of parameters served as a virtual participant. We then ran the experiment for each of these 90 virtual participants to generate simulated data. Last, we used hierarchical Bayesian modeling to fit these simulated data. Correlations between the estimated and the true parameters were calculated as metrics for model recovery. The correlations for the five parameters of the EWMV-VS model ranged from 0.79–0.89 (Figure S1), indicating highly accurate parameter estimation (2).

#### Supplemental effects-on-parameters analysis: controlling age and education year

The three group of participants had small but statistically significant differences in age and education year. To exclude the possibility that these demographic differences might contribute to the group differences in the decision characteristics assessed by the EWMV-VS model parameters, we performed supplemental analyses that include age and education year as the covariates. In particular, normalized age (or education year), along with NSSI, SDS and their interaction were entered into the effects-on-parameters analysis (EPA) as regressors. When age or education year was controlled, the group-level effects of NSSI, SDS and their interaction (Figures S2 and S3) were similar as those of the main EPA analysis (Figure 3 in the main text).

#### Supplemental statistical and modeling results comparing MDD Patients with NSSI<5 and NSSI≥5

To examine the effects of NSSI<5 versus NSSI≥5, we first performed an LMM analysis on $N_{pump}$ similarly as before, except that the regressor of participant group included four instead of three categories (see Methods). All the significant effects identified in the previous LMM were replicated in this new LMM (Figure 4B). That is, there were significant main effect of balloon type (*F*(1, 230.33)=231.00, *p<*0.001), trial number (*F*(1, 228.64)=50.97, *p<*0.001) as well as their interaction. The participant group also had a significant effect (*F*(3, 234.29)=2.65, *p=*0.049), but multi-comparison corrected *post-hoc* differences between the groups failed to reach significance.

Hierarchical Bayesian modeling analysis for four participant groups also suggests that the NSSI≥5 and NSSI<5 groups had similar model parameters (Figure 4C). Here we separately estimated the differences between D and HC, between the two NSSI groups and D, and between NSSI≥5 and NSSI<5 in each of the five parameters. Similar to what we reported earlier for NSSI as one group, participants with NSSI had higher choice consistency than the D group (difference *M*=0.24, 95% HDI [0.01,0.51]) and lower risk preference than the HC group (difference *M*=–0.11, 95% HDI [–2.1, –0.02]). The choice consistency of HC was also larger than D (difference *M*=0.19, 95% HDI [0.01,0.38]). No parameters showed significant differences between the NSSI≥5 and NSSI<5 groups (Supplemental Table S2).

#### Supplemental model comparison for 3-group and 4-group models

The results in the main text indicated that participants with NSSI<5 and NSSI≥5 had similar decision characteristics. Here we further compare the 3-group model (participants group: HC, MDD, D+NSSI) and 4-group model (participants group: HC, MDD, D+NSSI<5 , D+NSSI≥5) by separately fitting participants’ data using hierarchical Bayesian modeling method. As shown in Table S3, the 3-group model outperformed the 4-group, which provides direct support for the lack of differences between the D+NSSI<5 and D+NSSI≥5groups in decision characteristics.

### Supplemental tables

**Table S1. Expected log pointwise predictive density (ELPD) for the three models**

|  | **ELPD (LOO)** | **ΔELPD** | **weight** |
| --- | --- | --- | --- |
| **EWMV-VS** | –30572.67 | 0 | 0.77 |
| **EWMV** | –31134.89 | –563.07 | 0.18 |
| **Wallsten** | –31447.12 | –874.45 | 0.05 |

**Table S2. Parameter differences of the NSSI≥5 versus NSSI<5 groups**

|  | **Mean** | **95% HDI** |
| --- | --- | --- |
| $\boldsymbol{\Delta\phi}$ | 0.002 | [-0.022, 0.025] |
| $\boldsymbol{\Delta\xi}$ | 0.098 | [-0.241, 0.387] |
| $\boldsymbol{\Delta\beta}$ | -0.009 | [-0.495, 0.441] |
| $\boldsymbol{\Delta\lambda}$ | 0.001 | [-0.066, 0.068] |
| $\boldsymbol{\Delta\rho}$ | 0.669 | [-0.585, 2.010] |

**Table S3. Expected log pointwise predictive density (ELPD) for the 3-group and 4-group models**

|  | **ELPD (LOO)** | **ΔELPD** | **weight** |
| --- | --- | --- | --- |
| **3-group** | –30569.74 | 0 | 0.83 |
| **4-group** | –30571.58 | –1.85 | 0.17 |

### Supplemental figures

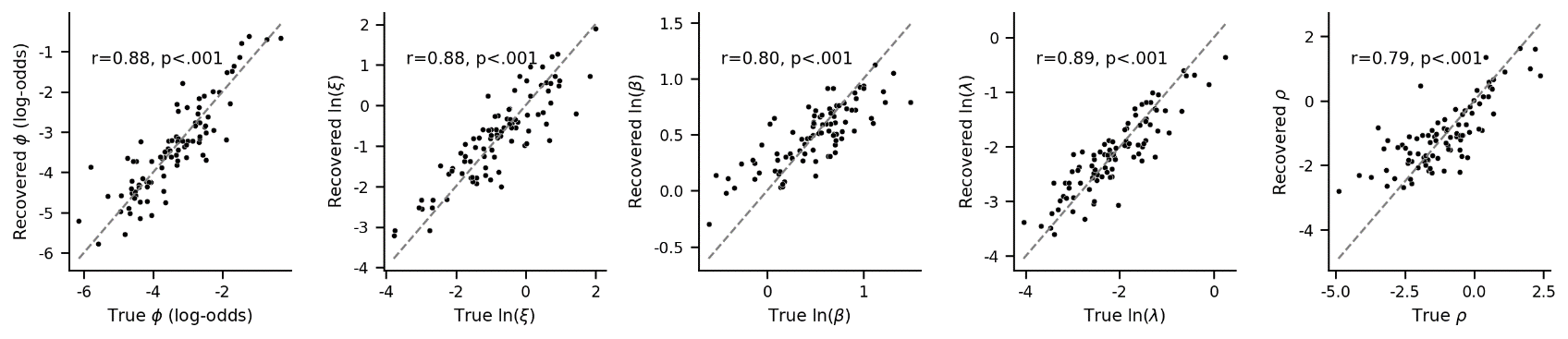

**Figure S1.** **Model recovery results of EVMV-VS model.** The parameters estimated from the simulated data (“recovered”) are plotted against the parameters used to generate the simulated data (“true”). The dashed line denotes the identity reference.

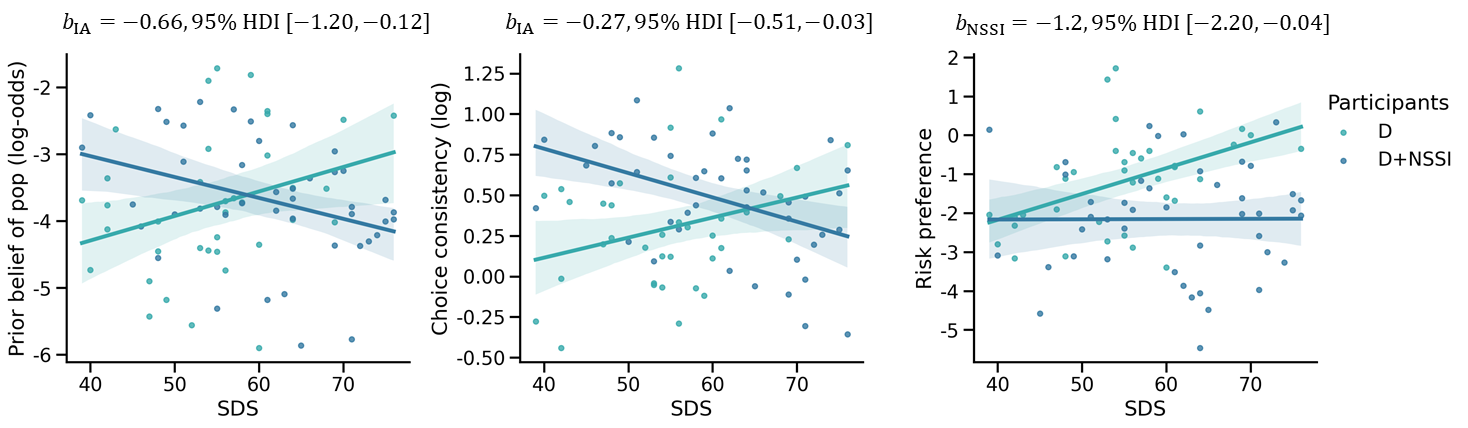

**Figure S2.** **Effects-on-Parameters analysis: controlling age.** Joint effects of NSSI and depression severity (as measured by the Self-Rating Depression Scale, SDS) on model parameters, with age as a covariate. Participants were limited to the D and D+NSSI groups with an overlapping range of SDS. Each dot denotes one individual participant. The shading on the regression line denotes the 95% HDI.

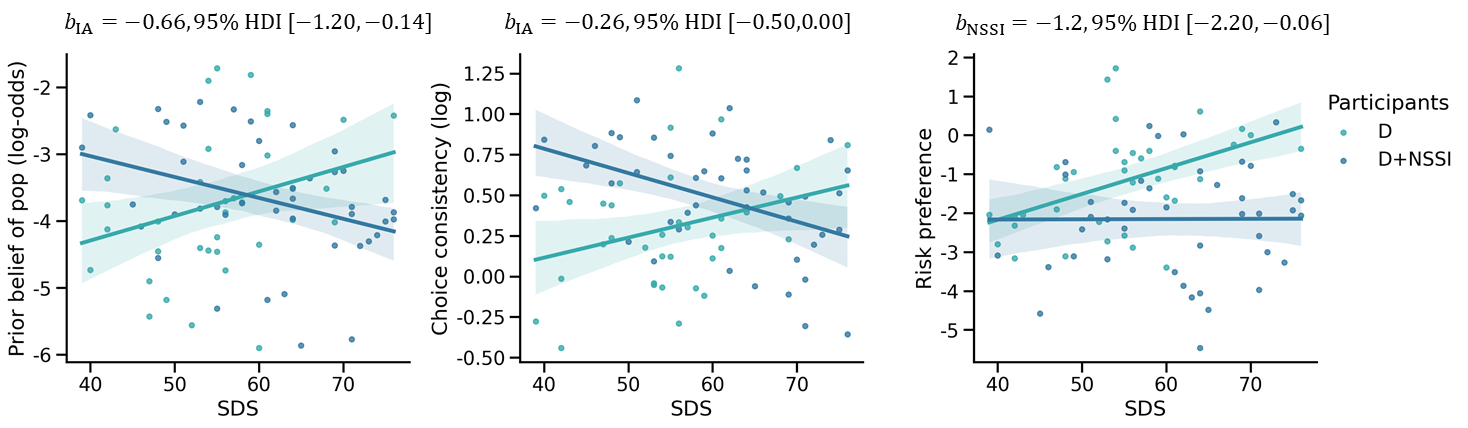

**Figure S3.** **Effects-on-Parameters analysis: controlling education year.** Joint effects of NSSI and depression severity (as measured by the Self-Rating Depression Scale, SDS) on model parameters, with education year as a covariate. Participants were limited to the D and D+NSSI groups with an overlapping range of SDS. Each dot denotes one individual participant. The shading on the regression line denotes the 95% HDI.
